# Toward machine-guided design of proteins

**DOI:** 10.1101/337154

**Authors:** Surojit Biswas, Gleb Kuznetsov, Pierce J. Ogden, Nicholas J. Conway, Ryan P. Adams, George M. Church

**Author notes:** Equal contribution (Surojit Biswas, Gleb Kuznetsov, Pierce J. Ogden).

## Abstract

Proteins—molecular machines that underpin all biological life—are of significant therapeutic and industrial value. Directed evolution is a high-throughput experimental approach for improving protein function, but has difficulty escaping local maxima in the fitness landscape. Here, we investigate how supervised learning in a closed loop with DNA synthesis and high-throughput screening can be used to improve protein design. Using the green fluorescent protein (GFP) as an illustrative example, we demonstrate the opportunities and challenges of generating training datasets conducive to selecting strongly generalizing models. With prospectively designed wet lab experiments, we then validate that these models can generalize to unseen regions of the fitness landscape, even when constrained to explore combinations of non-trivial mutations. Taken together, this suggests a hybrid optimization strategy for protein design in which a predictive model is used to explore difficult-to-access but promising regions of the fitness landscape that directed evolution can then exploit at scale.

## 1. Introduction

Proteins play critical roles in cells, including catalyzing reactions, sensing, signaling, and providing structure. Consequently, engineered proteins are of high therapeutic and industrial value, with a growing number of applications as medicines and industrial catalysts. A protein’s function is determined by its 3D structure, which is formed from a folded chain of amino acids. The challenge for protein engineers is to discover the sequence of amino acids that encodes a protein with desired function.

Despite advances in *in silico* prediction of protein folding, creating novel proteins from first principles remains limited to simple folds and static structures [1]. Instead, most protein engineers focus on improving existing proteins through directed evolution, structure-guided mutagenesis, and screening. Optimizing a protein’s function can be described as seeking the tallest peak in its fitness landscape [2], or searching the space of possible sequences for optimal function. Directed evolution is adept at searching a local neighborhood and climbing nearby peaks [3], but other functional neighborhoods may be separated by thin ridges or valleys that are only accessible by evolution over millions of years [4]. Therefore, a model with a sufficiently good approximation of the fitness landscape may guide the discovery of rare, high-functioning sequences that are otherwise inaccessible to directed evolution.

In this work, we explore the potential and challenges of closing the loop between machine learning and high-throughput wet lab experimentation to navigate a protein’s fitness landscape and ultimately optimize protein function (Figure 1). We demonstrate our ideas empirically using the green fluorescent protein (GFP) (Figure 2) as a representative test case, where we define the fitness of a specific GFP amino acid sequence to be its fluorescence—an easily measurable function.

**Figure 1:**
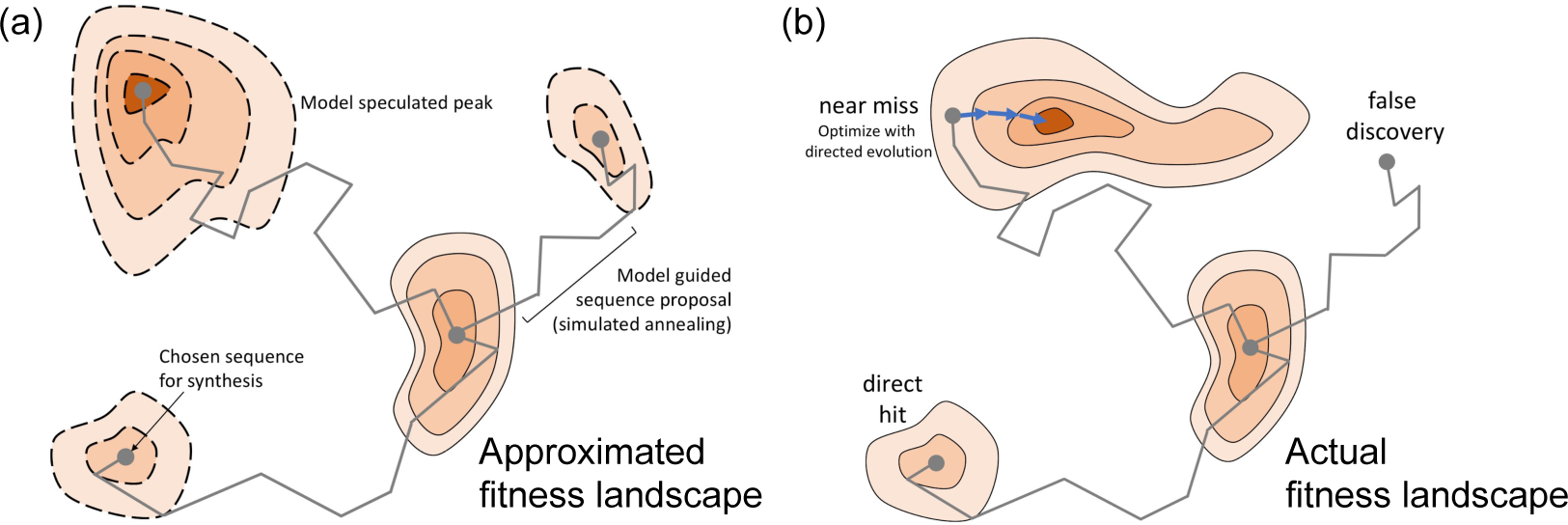
Workflow for ML-guided protein design. a) Simulated annealing of a model approximated fitness landscape proposes candidate sequences for exploration beyond the reach of directed evolution. b) The true landscape is iteratively revealed through synthesis and testing of proposed sequences. Model-guided designs can either be false discoveries, optimal, or near-misses (functional but not optimal). Directed evolution reveals the local neighborhood of model-guided designs and optimizes near misses.

**Figure 2:**
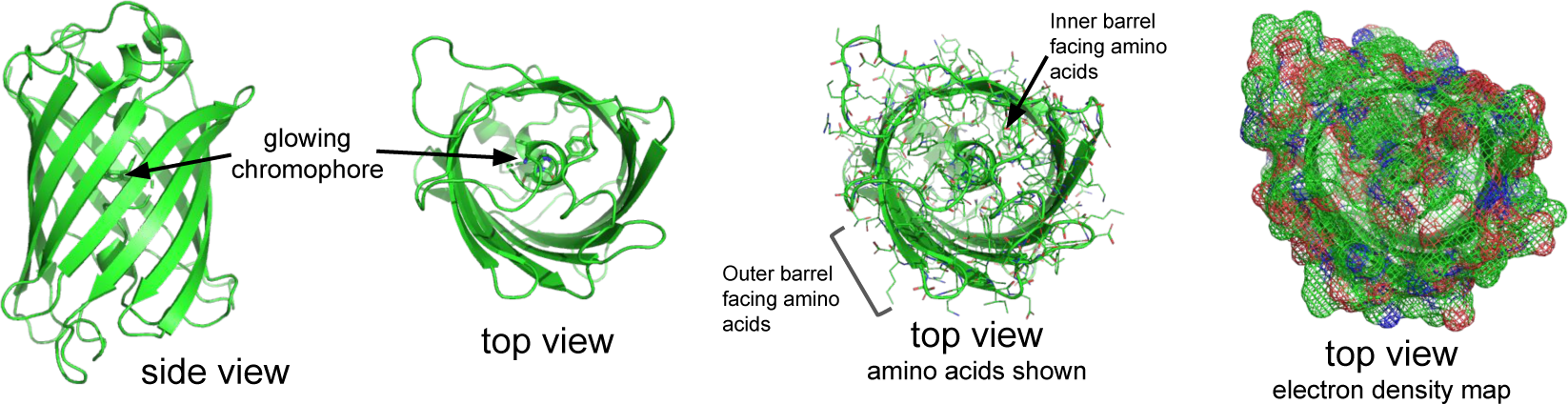
Structure of GFP. Left) side and top views of the GFP backbone illustrated as a “ribbon.” Right) Top views of GFP illustrating the orientation and interaction of amino acids inside and outside of the barrel. The electron density map better illustrates the natural occupancy of the amino acids.

In particular, we make the following contributions:

1. We investigate the challenges of selecting models for protein design that can generalize to unseen parts of the fitness landscape. Specifically, we first perform wet lab experiments to generate new, diverse datasets that we use to design several train-dev-test splits to compare models. We find that a novel, but simple model architecture we call *Composite Residues* performs best with respect to our generalization objectives.
2. We demonstrate the Composite Residues model’s ability to generalize using prospectively designed wet lab experiments. These experiments confirm the model’s ability to generalize to unseen parts of the landscape not only in terms of total-mutations-made, but also in terms of its ability to access difficult-to-reach regions of the landscape.
3. In an analysis of all wet lab validated gain-of-function mutants, including those obtained from a control directed evolution experiment, we demonstrate that non-trivial mutations— mutations that are hard to make while preserving function—have a high variance impact on protein function. Thus, while on average they negatively impact function, a significant minority of non-trivial mutations are predicted to significantly boost function.
4. We show with experiments that optimizing protein function via predictive modeling cannot easily “beat” directed evolution when given the same set of input sequences to optimize.

Taken together, we posit that a good optimization strategy for finding the highest functioning sequences may be to use supervised predictive models to explore difficult-to-reach, but functional sequence neighborhoods that directed evolution can then exploit in parallel and at scale (Figure 1).

## 2. Biological Background

Proteins are composed of chains of 20 naturally occurring amino acids, which are in turn encoded as a sequence of DNA. The protein’s biological function is a direct consequence of its structure, and therefore ultimately its sequence. How a change, or mutation, in the amino acid sequence encoding a protein affects its function is determined by which amino acid is being mutated as well as its adjacent amino acids in 3D space. Consequently, combinations of mutations may affect function in non-linear ways (epistasis), resulting in a complex, multi-modal fitness landscape that is difficult to optimize. Moreover, even for a small 100 amino acid protein the sequence space is immense (20^100^), and the vast majority of sequences are non-functional [5].

To explore a protein’s fitness landscape, we can leverage various molecular biology techniques to generate, read, and characterize the function of proteins by manipulating their respective coding DNA sequence. However, technological trade-offs in specificity and cost constrain the scope of sequences that can be characterized in any one experiment. Gene-length DNA synthesis remains expensive and limited to ≈100-1000 sequences per round of experiments. However, random mutagenesis (e.g., by error-prone PCR) of a given template DNA sequence is inexpensive and allows generating a library of DNA sequences in the local neighborhood of the template. Protein function can be measured in parallel, high-throughput assays where individual DNA sequences are physically linked to their function.

In the case of GFP, functional variants of the protein allow the cells to “glow.” Cells with differentially functional GFPs can be separated using Fluorescence-Activated Cell Sorting (FACS) combined with next-generation sequencing to characterize libraries of *≈* 10^5^ sequence-function pairs [6, 7].

In addition to a high-throughput assay, GFP offers several natural, intuitive controls to aid development of model-guided protein engineering methods. The protein consists of a barrel-shaped structure protecting a chromophore formed by three amino acids (Figure 2). The excitation-emission properties of GFP are directly affected by the identity of chromophore and inner-barrel amino acids, but are ultimately subject to the global stability determined by all 238 amino acids [8].

## 3. Related Work

Several studies have applied machine learning to predicting protein *structure* from sequence (e.g. [9, 10]). However, our work is focused on learning models that predict protein *function* directly from sequence so that we can use them to guide wet lab experiments toward a design goal [11].

Previous work leveraging machine learning methods to engineer proteins focused on variations of linear regression [12, 13], Gaussian process (GP) regression [14, 15], and other kernel-based methods to model the relationship between sequence and fitness to produce improved proteins.

Romero *et al.* demonstrated the utility of GP regression models for optimizing protein function [14]. This was effective for leveraging their training set of hundreds of protein variants; however, exact inference with GPs is impractical for applications beyond a few thousand training points [16]. Furthermore, kernel-based methods are fundamentally interpolative and are thus limited in their ability to explore beyond training data.

Fox *et al.* detailed an iterative, model-guided method for improving the function of an enzyme [13]. The authors performed 18 rounds of generating diversity, screening, and modeling to iteratively accumulate mutations that contributed to improved enzyme function. Despite using a purely additive model, Fox *et al.* observed that the combined effect of the final 35 mutations was not predicted by an additive model. By refitting a linear model only to the latest round of experiments, they were able to compensate for the simplicity of their model and traverse global non-linearity in the fitness landscape.

Learning models that can represent non-linear interactions may decrease engineering times and help to uncover proteins with unprecedented performance. Additionally, by pairing recent advances in high-throughput DNA synthesis and sequencing with multiplexed molecular biology, it is now possible to obtain the necessary quality and volume of data for principled model development. To our knowledge, no previous work on model-guided protein engineering has explored how to design and generate datasets that best complement modeling. In this work, we use high-throughput experimental capabilities not only to train models, but also to investigate when and why they generalize and how that ultimately affects engineering efforts.

## 4. Results

### 4.1 Experimental design and generation of datasets to guide model selection

In order to train and develop models for the purpose of protein design, we sought data that would help us assess model generalization. We employed wet lab techniques—gene-length DNA synthesis and error-prone PCR—to create datasets of functional GFP variants diverse in sequence and fluorescence intensity, complementing an already published local fitness landscape dataset of avGFP [7]. Our combined dataset of over 600,000 sequence-function pairs covered several distinct and previously unexplored functional neighborhoods of GFP (Supp. Figure 1). These initial training datasets are summarized as follows:

1. **Sarkisyan (avGFP)** - A set of ≈54,000 fluorescence measurements of randomly mutated variants of avGFP. These variants contained an average of 3 mutations and serve as a high-quality measurement of the local fitness landscape, or neighborhood, of avGFP.
2. **Break-Fix (sfGFP)** - sfGFP is a variant of avGFP differing by 14 mutations that are the result of several directed evolution projects to improve avGFP’s brightness and stability [17]. To complement the Sarkisyan dataset, we performed an exploration of the fitness landscape around sfGFP. Specifically, we employed an experimental strategy we refer to as Break-Fix Evolution to enrich for diverse mutants that maintain function by passing iteratively mutagenized variants through toggled rounds of positive and negative selection (Supp. Info. Section 2). We measured the brightness of ≈ 300,000 sequences quantitatively at coarse resolution (Supp. Info. Sections 3 and 4).
3. **96 Designed Neighborhoods** - The above two datasets represent thorough explorations of two GFP neighborhoods. To explore more neighborhoods in parallel, we proposed and then experimentally synthesized 96 “parent” sequences (5–40 mutations relative to sfGFP) using simulated annealing of a model approximated fitness landscape (Supp. Info. Section 5). To build this approximation we trained a fully-connected feed-forward neural network on the Sarkisyan and Break-Fix data based on the reasoning that the model could learn non-linear interactions among amino acids. We synthesized and characterized the 96 sequences in the wet lab and found that 43 out of 96 were functional. We then performed local mutagenesis (Supp. Info. Section 4) of these parents to generate neighborhoods and used Flow-Seq (Supp. Info. Section 3) to measure *≈*200,000 sequence-function pairs.

#### 4.1.1 Model Architectures

With the complete training set, diverse in function and edit distance, we proceeded to empirical model comparison. The proteins were featurized via their amino acid sequence. Each of the 20 amino acids was represented as a vector in ℝ^15^, which can be thought of as a character embedding for each element of a 238-character string. After flattening this sequences of embeddings, the protein representation was a 3,570-dimensional vector. We considered three architectures in support of our design goals. In each case, the amino acid embeddings were jointly learned with the model parameters.

1. **Linear Regression (LR)** - Flattened character embedded sequence passed as input to linear regression. ≈ 3,900 parameters.
2. **Feed Forward Network (FFN)** - Flattened character embedded sequence passed as input to a three hidden layer fully connected network with hidden layer dimensions of 100, 30, and 10. The network employed SELU [18] activations on all nodes, except the output. ≈ 361,000 parameters.
3. **Composite Residues (CR)** - We developed the Composite Residues architecture from the intuition that small, hierarchically organized groupings of interacting amino acids drive protein function. As such, in this architecture the sequence-length x 15 output matrix of character embedding is passed through a Composite Residues layer, in which the embedded sequence is multiplied from the left by a 5 x sequence-length pooling matrix. The rows of the resulting 5 ×; 15 compressed sequence represent “composite” amino acid residues. This matrix is then flattened and passed to a 5-dimensional fully connected hidden layer with SELU activations that interprets the learned groupings and feeds an output node. Further details can be found in Section 6 of the Supplemental Information. ≈1,900 parameters.

For each model class, the specific architectures described above were arrived upon through several empirical iterations of hyperparameter tuning and model assessment using Sarkisyan as a training set and Break-Fix as a test set, and vice versa. For each architecture, we also compared the effect of having a linear versus sigmoid activation on the final output node. The models were trained using mean-squared error (MSE) in log relative fluorescence units.

#### 4.1.2 Model selection is sensitive to train-dev-test split and performance metric

Proper development (“dev”) and test datasets as well as choice of performance metrics are key factors for arriving at useful models. A good development dataset—an out-of-sample dataset used to benchmark the model during training—should overlap in distribution not only with the training data, but also with the target test cases to which we hope to generalize. Models can only be directly compared within a train-dev-test dataset split and model selection should be based performance on dataset splits that are most consistent with the design (generalization) objective.

In the protein design domain, the characteristics of an effective dev set have not been previously explored. We enumerated several natural characteristics to consider: edit distance to a reference point (e.g., wild-type avGFP), phenotype distribution (e.g., dark vs bright), and positional distribution of amino acids.

With these considerations in mind, we created 4 different train-dev-test splits:

1. **Sarkisyan 85-5-10** - Random split of the Sarkisyan dataset with 85% train, 5% dev, and 10% test.
2. **Sarkisyan-Break-Fix** - Train and dev data from Sarkisyan (random 90% train, 10% dev). Test data from Break-Fix.
3. **96-Designs-Sarkisyan** - Train-dev on the 96 Designed Neighborhoods data (random 90% train, 10% dev). Test on Sarkisyan.
4. **96-Designs-Kamchatka-Holdout** - Train on 96 Designed Neighborhoods data with one far neighborhood (15 mutations from sfGFP and most other parents) held out as the test set, which we nicknamed “Kamchatka” based on its spatial location in a 2D PCA projection of our dataset (Figure 3, upper-right panel). We also synthesized an “ancestral” sequence of Kamchatka that had 3 of its founding mutations and measured its neighborhood. We used this ancestral neighborhood as the dev set.

**Figure 3:**
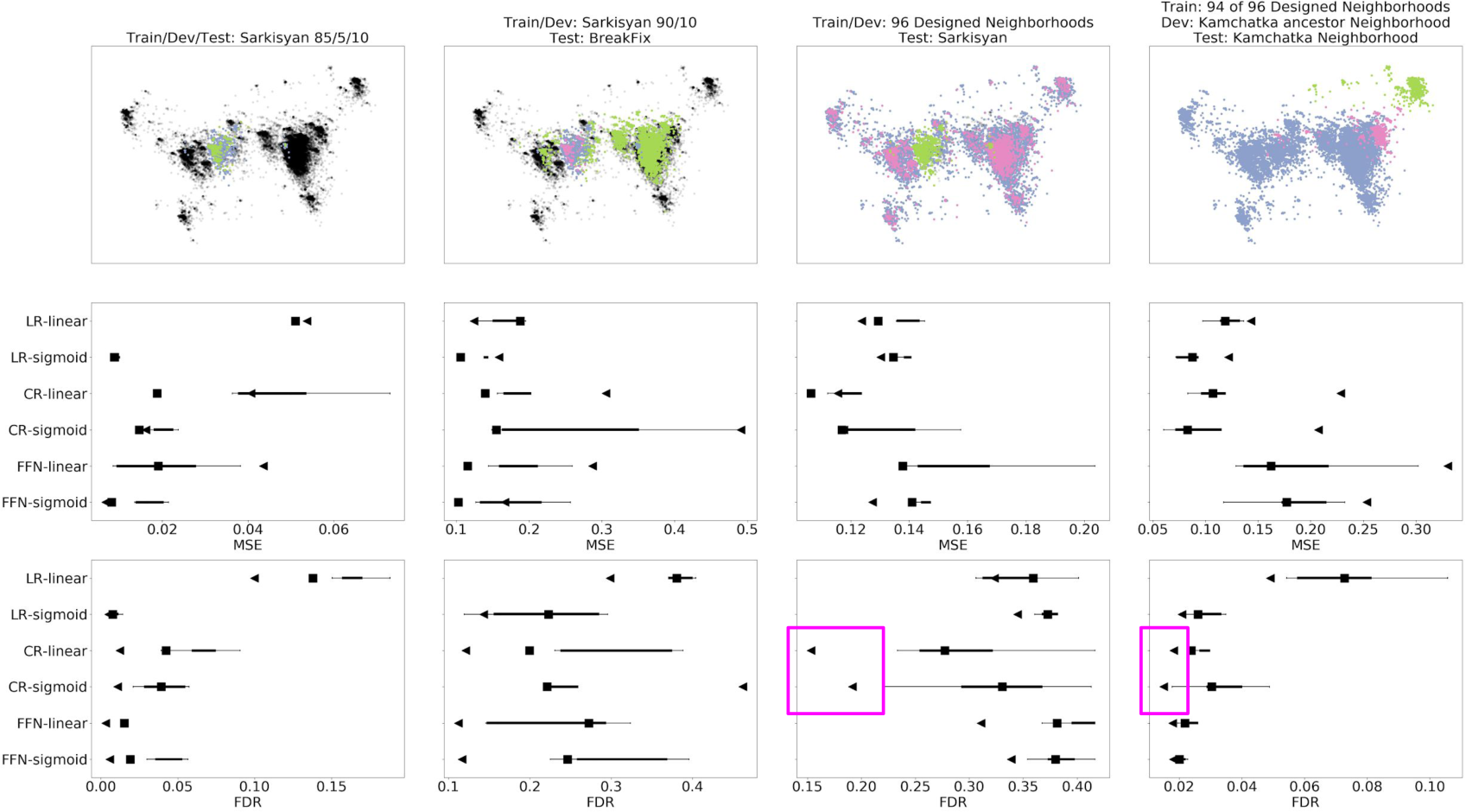
Variations of train-dev-test split design guide model selection. (Top) The three experimentally-generated datasets are visualized using the first two principal components of a PCA (see also Supp. Figure 1). Top panels indicate distribution of train (blue), dev (pink), test (green) data points. Black points are all remaining, unused points. (Bottom) Four train-dev-test splits were chosen for comparing six model architectures across two performance criteria, MSE and FDR. Box plots indicate quartile boundaries of replicate performance. Triangle and square points indicate the min- and mean-ensembled performance of the replicates, respectively.

We trained the models described in the previous section, and performed comparisons within each train-dev-test split (Figure 3, Supp. Info. Section 7). In addition to the mean squared error (MSE), we computed the false discovery rate (FDR) on the held-out test set using a reasonable binarization threshold. FDR is a critical metric for protein design as it is directly related to money and effort lost on non-functional sequences.

We performed five training replicates of each model by using a different random weight initialization each time. For a given model architecture, we built min- and mean-ensembles of the replicates to test their potential for learning different, but complementary distributions. Here, for example, a min-ensemble would return the minimum predicted value of all replicate models for a given input.

The train-test-split analysis in Figure 3 illustrates the challenges of identifying proper dev and test sets for protein engineering. In the naïve random train-test-split (Sarkisyan 85-5-10), a linear model with sigmoid activation (LR-sigmoid) strongly out-performed the other models. In fact, LR-sigmoid performed reasonably well on all datasets considered. However, among the more non-standard train-test-split designs employing the 96 Designed Neighborhoods, we found the Composite Residues architecture with a sigmoid activation (CR-sigmoid) to perform comparably to LR-sigmoid according to MSE, and out-perform LR-sigmoid according to FDR. This was especially true of the Composite Residues min-ensemble. However, on 96 Designed Neighborhoods with Kamchatka hold-out, even the FFN model was competitive with CR-sigmoid, emphasizing the challenge of model selection for protein design.

With the goal of discovering new, distant variants of GFP that remain functional, we considered FDR to be more important as it controls the amount of experimental capital used on non-functional sequences. We selected the CR-sigmoid model to continue our explorations as it performed well on train-test-splits most in line with our generalization objectives. Additionally, we found that the Composite Residues model weights offered intuitive biophysical interpretation, which assured us further of the model’s ability to generalize (Supp. Info. Section 6).

### 4.2. Models trained on local fitness data can generalize non-locally

We next designed wet lab experiments to test the ability of the min-ensembled Composite Residues model to generalize to unseen parts of the fitness landscape.

#### Unrestricted exploration of the fitness landscape

– Examining the brightness of the 96 Designed Neighborhoods parents as a function of number-of-mutations revealed that on average 20+ mutations could be made to the protein without completely impairing function (Figure 4a, green circles). By contrast, random mutagenesis—an experimental baseline—typically destroyed function after five mutations ([7], and our sfGFP Break-Fix data). This suggested that even the non-optimal FFN model could generalize to unseen parts of the fitness landscape.

**Figure 4:**
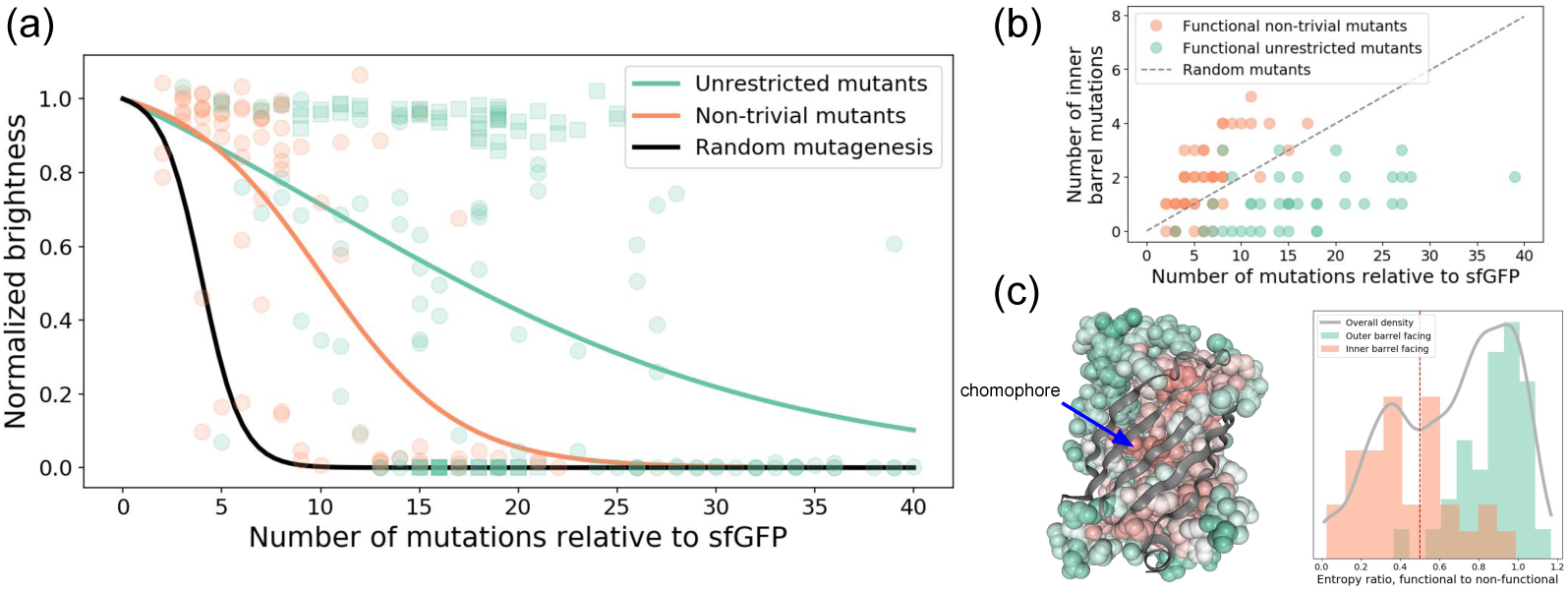
Locally trained Composite Residues model can generalize to non-local parts of the landscape. (a) Normalized brightness (log-scale) vs number of mutations for experimentally generated random mutants (black), non-trivial mutants designed by CR-sigmoid (orange), and unrestricted mutants (green). Green circles indicate parents of initial 96 Designed Neighborhoods. Green squares correspond to unrestricted mutants designed using the Composite Residues model starting from the bright 96 Designed Neighborhoods parents. (b) Inner-barrel facing mutations are enriched in functional non-trivial mutants relative unrestricted mutants. Gray dashed line shows the expected number of inner-barrel mutations assuming random mutagenesis. (c) Atomic space-filled structure of GFP with core exposed and amino acids colored according to positional entropy ratios. Right plot shows the distribution of positional entropy ratios (functional-to-nonfunctional) for all amino acids (grey line), inner-barrel facing (orange), and outer barrel facing (green). Red vertical line depicts a hard threshold for defining non-triviality.

We next experimentally verified that the min-ensembled Composite Residues model could likewise move through the landscape when required to make at least 15 mutations to one of the 96 Designed Neighborhoods parent sequences via unrestricted simulated annealing (Supp. Info. Section 5). We found that in testing these proposed unrestricted mutants, the CR model not only achieved this objective, but also improved upon or maintained brightness near wild-type sfGFP (Figure 4a, green squares).

Examining the properties of these designed sequences in the context of structure, we noticed functional mutants were enriched for outer-barrel mutations (Figure 4b). Even though the model was not provided with the structure, it was able to learn that outer-barrel amino acids tended to be more mutable without impairing function (matching literature and intuition). Thus, perhaps one trivial way to generalize to unseen parts of the fitness landscape might simply be to progressively mutate permissible amino acids.

#### Restricted exploration of the fitness landscape - generalization to non-trivial mutants

– To test whether the min-ensembled CR model could explore more difficult-to-reach regions of the landscape, we defined a measure of how non-trivial each amino acid is to mutate based on local mutagenesis data. Specifically, we calculated the Shannon entropy at each sequence position of all bright sequences and divided it point-wise by the Shannon entropy at each position of all dark sequences. This entropy ratio should be closer to zero for immutable amino acids, and closer to one for permissible amino acids. Indeed, inner-barrel amino acids tended to have lower entropy ratios than outer-barrel ones (Figure 4c).

We used these non-triviality scores to bias simulated annealing sampling of the CR model and proposed 76 new GFP sequences enriched for non-trivial mutations (Supp. Info. Section 5, Figure 4b). Wet lab validation confirmed these model-designed non-trivial mutants maintained function better than random mutagenesis (Figure 4a).

### 4.3 Non-trivial generalization may be critical for protein design

We next investigated how the above lessons and ways of thinking about generalization might impact protein design. In parallel to evaluating designed sequences, we performed two rounds of (wet lab) directed evolution, selecting for brighter sequences in each round (Supp. Info. Section 8). Directed evolution produced a number of gain-of-function (GoF; brighter than sfGFP) mutants. Several ML-proposed designs were of comparable brightness, but were not statistically brighter (Supp. Figure 5).

**Figure 5:**
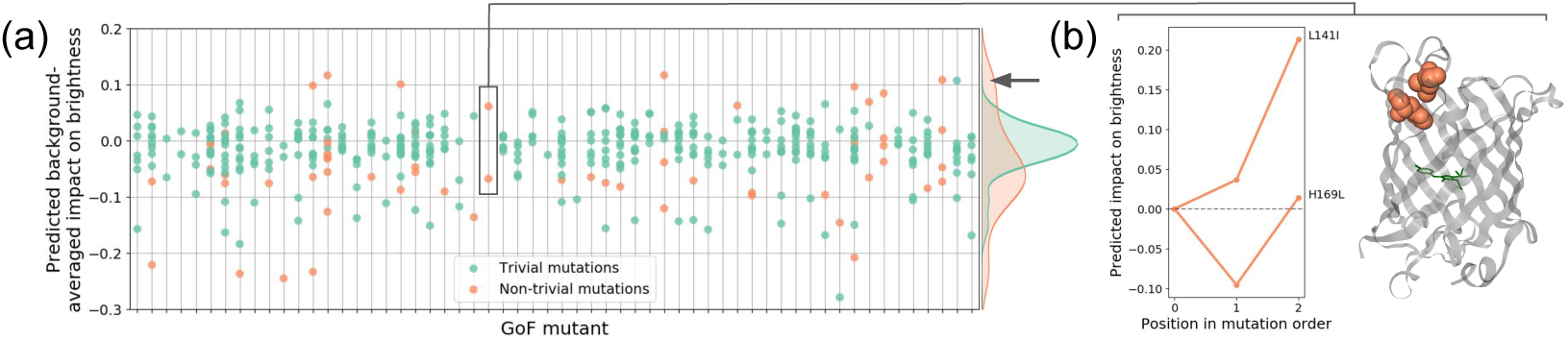
Non-trivial mutations exhibit higher variance effect on function. (a) Predicted background-averaged impact on brightness for all mutations in all brighter-than-sfGFP mutants. (b) Example of non-additive effect of two mutations learned by the model and visualized on GFP structure.

We examined the mutational composition of these GoF sequences. With a few exceptions, both designed and evolved sequences tended to have more trivial than non-trivial mutations. Using the CR model, we computed the predicted background-averaged impact on brightness for each mutation within a sequence (Figure 5a, Supp. Info. Section 10). We observed that trivial mutations tended to have small effect sizes hovering around neutral. Suprisingly, however, we noticed that even though non-triviality was defined on a loss-of-function basis (i.e. mutations at a non-trivial position tended to break protein function on average), a significant minority of non-trivial mutations were predicted to have a strongly positive impact on brightness (Figure 5a).

One GoF mutant had only two non-trivial mutations, one with positive background-averaged impact (sequence position 141) and the other with negative background-averaged impact (position 169). We examined the predicted effect of each mutation whether it occurs alone or in the context of the other mutation (Figure 5b, left). This revealed a predicted non-linearity in which the individual impact of the mutation at 169 was negative without the mutation at 141, but was neutral or positive when 141 was in the background. Similarly, in the context of the 169 mutation, the predicted impact of the mutation at 141 was significantly enhanced (i.e. positive epistasis). Further, we found that amino acids 141 and 169 are in close physical proximity (Figure 5b, right). Moreover, they co-localize at the lid of the GFP barrel, which is a likely folding nucleus important for proper GFP maturation [19].

We conclude that though non-trivial mutations tend to have a negative phenotypic impact, they induce greater phenotypic variance and thus offer opportunity for making relatively large improvements in function with only a few mutations. This suggests pursuing non-trivial mutations aggressively in terms of training dataset design, modeling effort, and sequence proposal may be valuable for protein design.

## 5 Discussion

Our study has practical implications for how to effectively combine high-throughput molecular biology and machine-learning for protein engineering. At first glance, directed evolution appears to be a wet lab baseline to challenge with a machine-guided approach. While directed evolution only performs local optimization of the fitness landscape, it achieves unparalleled scale and performs local exploitation well because it can screen millions of mutants per evolution cycle. Nevertheless, we observed that an appropriately selected model can perform exploration by making non-trivial leaps into the fitness landscape that directed evolution alone cannot do. Taken together, this suggests a hybrid optimization strategy in which, per design cycle, a model is used to non-locally explore promising regions of the fitness landscape that directed evolution can then optimize and exploit.

Given model-guided exploration of the fitness landscape may be a valuable goal, we showed several ways in which we could think about the generalization of our models. The simplest demonstration of generalization is to show that a model can effectively propose functional protein sequences that are many mutations away from a reference point and locally associated training data. A more difficult demonstration is to show the same model can effectively propose functional protein sequences that contain many non-trivial mutations. We observed that, perhaps due to their extensive structural involvement, non-trivial amino acids may be points of high leverage with respect to function optimization. We speculate that this may be because such mutations fundamentally change or reconfigure the protein, thereby enabling access to different and possibly higher functional optima.

Of course, the discussion so far assumes there is a model in hand with which to assess generalization. A key contribution of this work is to highlight the difficulty of selecting a strongly generalizing model in the protein engineering domain. We found that model selection depended heavily on train-dev-test split design. Interestingly, the model with the fewest number of parameters, Composite Residues, was the one that generalized the best in our evaluations. This, in part, is a consequence of the relatively simple out-of-sample hypotheses the model can make. However, we also argue that the superior performance of Composite Residues is attributable to its architecture, which succinctly captured realistic modularity in how groups of amino acids might biophysically cooperate to impact protein function.

While we have developed our proof-of-concept using GFP, we expect that the methods, design principles, and lessons learned extend broadly to other proteins. All proteins fold according to the same thermodynamic laws, and have variably mutable amino acids that offer differing amounts of leverage over the protein. From a machine learning perspective, this makes protein engineering an exciting domain for developing new methods as the effect of mutational combinations range from purely additive to highly non-linear. Importantly, unlike many other design domains (e.g., chemical design [20]), synthesizing protein designs is largely a digital exercise (i.e. ordering sequences of DNA), and testing them experimentally can often be done massively in parallel.

## 6 Acknowledgements

We thank Daniel B. Goodman and John Aach for valuable feedback on the manuscript. SB was supported by an NIH Training Grant to the Harvard Bioinformatics and Integrative Genomics program as well as a NSF GRFP Fellowship. GK was supported by an NIH Training Grant to the Harvard Biophysics Program. PJO was supported by the National Human Genome Research Institute (RM1HG008525). RPA was partially funded by NSF IIS-1421780 and the Alfred P. Sloan Foundation. Experimental work was supported by the U.S. Department of Energy, Office of Science, Office of Biological and Environmental Research under Award Number DE-FG02-02ER63445. Computational resources were generously provided by the AWS Cloud Credits for Research program.

## References

[1] Po-Ssu Huang, Scott E Boyken, and David Baker. The coming of age of de novo protein design. Nature, 537(7620):320–327, 2016.

[2] Sewall Wright. The roles of mutation, inbreeding, crossbreeding, and selection in evolution, volume 1. na, 1932.

[3] Philip A Romero and Frances H Arnold. Exploring protein fitness landscapes by directed evolution. Nat. Rev. Mol. Cell Biol., 10(12):866–876, 2009.

[4] Victoria Pokusaeva, Dinara Usmanova, Ekaterina Putintseva, Lorena Espinar, Karen Sarkisyan, Alexander Mishin, Natalya Bogatyreva, Dmitry Ivankov, Guillaume Filion, Lucas Carey, et al. Experimental assay of a fitness landscape on a macroevolutionary scale. bioRxiv, page 222778, 2018.

[5] J M Smith. Natural selection and the concept of a protein space. Nature, 225(5232):563–564, February 1970.

[6] Sriram Kosuri, Daniel B Goodman, Guillaume Cambray, Vivek K Mutalik, Yuan Gao, Adam P Arkin, Drew Endy, and George M Church. Composability of regulatory sequences controlling transcription and translation in Escherichia coli. Proc. Natl. Acad. Sci. U. S. A., 110(34):14024–14029, August 2013.

[7] Karen S Sarkisyan, Dmitry A Bolotin, Margarita V Meer, Dinara R Usmanova, Alexander S Mishin, George V Sharonov, Dmitry N Ivankov, Nina G Bozhanova, Mikhail S Baranov, Onuralp Soylemez, Natalya S Bogatyreva, Peter K Vlasov, Evgeny S Egorov, Maria D Logacheva, Alexey S Kondrashov, Dmitry M Chudakov, Ekaterina V Putintseva, Ilgar Z Mamedov, Dan S Tawfik, Konstantin A Lukyanov, and Fyodor A Kondrashov. Local fitness landscape of the green fluorescent protein. Nature, 533(7603):397–401, May 2016.

[8] Peter Dedecker, Frans C De Schryver, and Johan Hofkens. Fluorescent proteins: shine on, you crazy diamond. J. Am. Chem. Soc., 135(7):2387–2402, February 2013.

[9] Vladimir Golkov, Marcin J Skwark, Antonij Golkov, Alexey Dosovitskiy, Thomas Brox, Jens Meiler, and Daniel Cremers. Protein contact prediction from amino acid co-evolution using convolutional networks for graph-valued images. In Advances in Neural Information Processing Systems, pages 4222–4230, 2016.

[10] Mohammed AlQuraishi. End-to-end differentiable learning of protein structure. bioRxiv, page 265231, 2018.

[11] David A Cohn, Zoubin Ghahramani, and Michael I Jordan. Active learning with statistical models. Journal of artificial intelligence research, 4:129–145, 1996.

[12] Jun Liao, Manfred K Warmuth, Sridhar Govindarajan, Jon E Ness, Rebecca P Wang, Claes Gustafsson, and Jeremy Minshull. Engineering proteinase K using machine learning and synthetic genes. BMC Biotechnol., 7:16, March 2007.

[13] Richard J Fox, S Christopher Davis, Emily C Mundorff, Lisa M Newman, Vesna Gavrilovic, Steven K Ma, Loleta M Chung, Charlene Ching, Sarena Tam, Sheela Muley, John Grate, John Gruber, John C Whitman, Roger A Sheldon, and Gjalt W Huisman. Improving catalytic function by ProSAR-driven enzyme evolution. Nat. Biotechnol., 25(3):338–344, 2007.

[14] Philip A Romero, Andreas Krause, and Frances H Arnold. Navigating the protein fitness landscape with Gaussian processes. Proc. Natl. Acad. Sci. U. S. A., 110(3):E193–201, January 2013.

[15] Claire N Bedbrook, Kevin K Yang, Austin J Rice, Viviana Gradinaru, and Frances H Arnold. Machine learning to design integral membrane channelrhodopsins for efficient eukaryotic expression and plasma membrane localization. PLoS Comput. Biol., 13(10):e1005786, October 2017.

[16] Elad Gilboa, Yunus Saatçi, and John P Cunningham. Scaling multidimensional inference for structured Gaussian processes. IEEE Trans. Pattern Anal. Mach. Intell., September 2013.

[17] Jean-Denis Pédelacq, Stéphanie Cabantous, Timothy Tran, Thomas C Terwilliger, and Geoffrey S Waldo. Engineering and characterization of a superfolder green fluorescent protein. Nat. Biotechnol., 24(1):79–88, January 2006.

[18] Günter Klambauer, Thomas Unterthiner, Andreas Mayr, and Sepp Hochreiter. Self-normalizing neural networks. In Advances in Neural Information Processing Systems, pages 972–981, 2017.

[19] Matthew H Zimmer, Binsen Li, Ramza Shahid, Paola Peshkepija, and Marc Zimmer. Structural consequences of chromophore formation and exploration of conserved lid residues amongst naturally occurring fluorescent proteins. Chemical physics, 429:5–11, 2014.

[20] Rafael Gómez-Bombarelli, Jorge Aguilera-Iparraguirre, Timothy D Hirzel, David Duvenaud, Dougal Maclaurin, Martin A Blood-Forsythe, Hyun Sik Chae, Markus Einzinger, Dong-Gwang Ha, Tony Wu, et al. Design of efficient molecular organic light-emitting diodes by a high-throughput virtual screening and experimental approach. Nature materials, 15(10):1120, 2016.

